# Phenotypic Assessment of Clinical *Escherichia coli* Isolates Predicts Uropathogenic Potential

**DOI:** 10.1101/2022.08.31.506135

**Authors:** A.E. Shea, A.E. Frick-Cheng, S.N. Smith, H.L.T. Mobley

## Abstract

For women in the United States, urinary tract infections (UTI) are the most frequent diagnosis in emergency departments, comprising 21.3% of total visits. Uropathogenic *Escherichia coli* (UPEC) causes ∼80% of uncomplicated UTI. To combat this public health issue, it is vital to characterize UPEC strains as well as differentiate them from commensal strains to reduce the overuse of antibiotics. Surprisingly, no genetic signature has been identified which clearly separates UPEC from other *E. coli*. Therefore, we examined whether phenotypic data could be predictive of uropathogenic potential. We screened 13 clinical strains of UPEC, isolated from cases of uncomplicated UTI in young otherwise healthy women, in a series of microbiological phenotypic assays using UPEC prototype strain CFT073 and non-pathogenic *E. coli* strain MG1655 K12 as controls. Phenotypes included adherence, iron acquisition, biofilm formation, human serum resistance, motility, and stress resistance. These data were able to predict the severity of bacterial burden in both the urine and bladders using a well-established experimental mouse model of UTI. Multiple linear regression using three different phenotypic assays, growth in minimal medium, siderophore production, and type 1 fimbrial expression, was predictive of bladder colonization (adjusted r^2^=0.6411). Growth in *ex vivo* human urine, hemagglutination of red blood cells, and motility modeled urine colonization (adjusted r^2^=0.4821). These results showcase the utility of phenotypic characterization to predict the severity of infection these strains may cause. We predict that these methods will also be applicable to other complex, genetically redundant, pathogens.

**Importance:** Urinary tract infections are the second leading infectious disease worldwide, occurring in over half of the female population during their lifetime. Most infections are caused by uropathogenic *Escherichia coli* (UPEC). These strains can commensally colonize the gut, but upon introduction to the urinary tract, can infect the host and cause disease. Clinically, it would be beneficial to screen patient *E. coli* strains to understand their pathogenic potential, which may lead to the administration of prophylactic antibiotic treatment for those with increased risk. Others have proposed the use of PCR-based genetic screening methods to detect UPEC and differentiate them from other *E. coli* pathotypes; however, this method has not yielded a consistent uropathogenic signature. Here, we have used phenotypic characteristics such as growth rate, siderophore production, and expression of fimbriae to successfully predict uropathogenic potential.

## Introduction

Half of women will experience at least one urinary tract infection (UTI) in their lifetime (1). These ubiquitous infections cause five billion dollars in associated healthcare costs in the US and annually affect 150 million women worldwide (2, 3). While a variety of bacteria cause UTI in uncomplicated cases, the predominant etiological pathogen is uropathogenic *Escherichia coli* (UPEC) which causes over 80% of uncomplicated cases (4, 5). Therefore, to combat this public health issue, it is vital to fully understand and characterize individual UPEC strains and beneficial to differentiate UPEC from commensal *E. coli*.

UPEC is part of the extraintestinal pathogenic *E. coli* (ExPEC) grouping, which encompasses any *E. coli* that causes disease outside of the gut (6, 7). Unlike other *E. coli* pathotypes, which can be identified by specific virulence gene repertoires, UPEC has incredible genetic heterogeneity (8, 9) and encodes a wide variety of virulence factors: up to six virulence-associated iron acquisition systems, six toxins, and 13 different adhesins (3, 10-12). Some strains produce almost all these virulence factors, while others produce only a select few. One of the few conserved virulence factors is the highly studied type 1 fimbriae; however, commensal isolates such as K12 also encode this adhesin (13, 14), Furthermore, *E. coli* that cause asymptomatic bacteriuria (ABU) are genetically similar to strains that cause symptomatic infection (15, 16).

A few studies have attempted to develop a diagnostic PCR assay to differentiate between ExPEC and diarrheagenic strains (17, 18). One study showed if a strain encoded *fyuA, chuA, vat* and *yfcV* it was ten times more likely to be a UPEC or neonatal meningitis *E. coli* (NMEC) isolate (17). However, they were unable to differentiate between UPEC and NMEC. Moreover, only 58% of their UPEC cohort encoded *vat*, and 69% encoded *yfcV*, leaving a relatively large population of UPEC overlooked.

Another study sought to differentiate between non-invasive UPEC, (ABU or cystitis-causing) and invasive UPEC (pyelonephritis or bacteremia-causing) and found *papG2* was highly enriched in the invasive strains (19). Although an important finding, there is a need to differentiate between ABU and cystitis-causing strains, especially given current clinical guidelines state ABU should not be treated with antibiotics while cystitis is (20-22). Indeed, ABU strains have been proposed as a treatment to prevent symptomatic cystitis (23, 24). Therefore, there is clearly a need to distinguish ABU strains from those that elicit pathogenesis.

UPEC can also transiently colonize the gut or the periurethral region of the host (25, 26). This wide tissue tropism obfuscates which virulence factors are required to infect the urinary tract as opposed to factors used for these other niches and UPEC will encode functionally redundant genes. For example, iron acquisition is essential for host colonization and UPEC encodes up to four different siderophores and two different heme receptors (27-29) and deleting a single system is not sufficient to completely ablate fitness (29). UPEC strains typically encode any number of combinations of these systems; A particular virulence factor is not vital, but the combination is important to achieve uropathogenesis. Consequently, this makes UPEC identification via a single gene, or even several genes, difficult.

To address this, we hypothesized that measuring phenotypic experimental outcomes, rather than genotyping, will predict infectivity. We characterized 13 clinical UPEC isolates (30, 31), and compared their behavior to well-characterized UPEC type strain CFT073 and intestinal commensal isolate K12. We performed 18 *in vitro* phenotypic assays associated with known virulence mechanisms of UPEC and correlated the results with an experimental mouse model of UTI. We were able to model bacterial burden in both the urine and bladders via multiple linear regression using three different phenotypic assays each. Growth in minimal medium, siderophore production and type 1 fimbrial expression was predictive of bladder colonization (adjusted r^2^=0.6411) while growth in *ex vivo* human urine, hemagglutination of red blood cells and motility modeled urine colonization (adjusted r^2^=0.4821). These results indicate the utility of phenotypic characterization to reduce the complexity of diverse, redundant genomes of UPEC to predict severity of infection.

## Results

### UPEC display diverse sugar metabolism efficiencies and iron acquisition abilities

Rapid growth and metabolic flexibility are key for UPEC survival and infection (31-33). Therefore we monitored the metabolic switch from anaerobic to aerobic conditions using phenol red as a pH indicator to model the switch from the gut to bladder environment. Both anaerobic respiration and fermentation result in lactic acid secretion thus lowering the local pH and turning the bacterial colony yellow. Aerobic respiration leads to the production of carbon dioxide and water, yielding a more red/pink color for the bacteria. Under both aerobic and anaerobic conditions, the clinical isolates were tested on each of four carbohydrates (glucose, glycerol, galactose, or ribose) as a preferred primary carbon source in LB to mimic anaerobic fermentative conditions (**Fig. 1A**). We observed that on LB plates under aerobic conditions, all strains performed aerobic respiration (**Fig. 1A**). Under anaerobic conditions all strains performed fermentation but quickly switched to aerobic respiration after 6h in an aerobic environment (**Fig. 1B**) However, introduction of carbohydrates delayed the switch to aerobic respiration (**Fig. 1B**) and likely encouraged utilization of glycolysis and the pentose phosphate pathway. Only when the carbohydrate was depleted, would bacteria switch over to exclusively amino acid-based metabolic processes as evidenced by a deepening shift from orange to red only after 24 hours of post-aerobic incubation.

**FIG 1.**
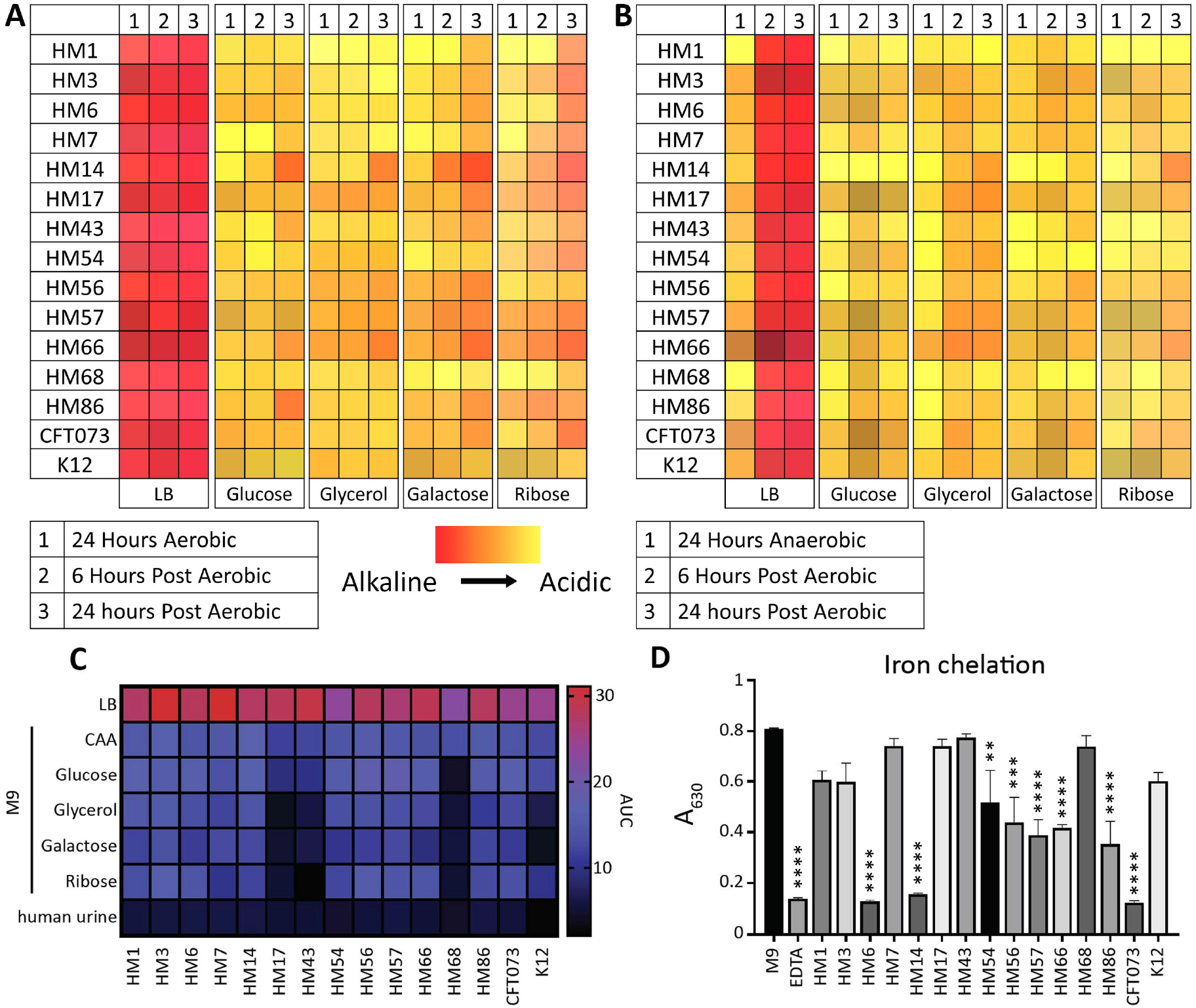
Three UPEC isolates display growth defects on sugar carbon sources and minimal chelation ability. UPEC isolates were spotted from overnight cultures onto LB agar plates containing different indicated carbon sources and 0.1% phenol red pH indicator. Plates were initially incubated at 37°C in either (A) aerobic or (B) anaerobic conditions. Plates were then transitioned to the benchtop for additional time point observation. The colors represent the hues displayed on the agar, with more red being more alkaline and yellow indicating acidic end products. (C) UPEC strains were inoculated 1:100 from overnight cultures into different media types. Strains were incubated at 37°C with aeration for 24 h and OD_600_ was recorded every 15 min. AUC was calculated and visually represented. Data displayed are a mean of six biological replicates. (D) Strains were cultured overnight, shaking at 37°C in M9 minimal medium supplemented with 0.4% glucose. 100 µL of supernatants was combined with 100 µL of chrome azurol S (CAS) shuttle solution and incubated at room temperature for 30 m. 10 mM EDTA served as a positive control and M9 medium only served as a negative control. After 30 minutes, the absorbance at 630 nM was read. Data are displayed as the mean of four biological replicates, error bars indicate SEM. One-way ANOVA was performed using with Dunnet’s multiple test corrections compared to M9. **, *P* < 0.01; ***, *P* < 0.005; ****, *P* < 0.0001.

Our previous study (34) has shown that global gene expression in LB recapitulates the transcriptome during human UTI. Pooled human urine can also recapitulate a similar expression profile for specific subsets of genes, such as those that control iron acquisition. Our lab has also found that UPEC prefers amino acids as a carbon source in both human and murine UTI (31, 34, 35). We wanted to investigate whether there were any strain-specific differences in growth when using carbohydrates as a sole carbon source as opposed to amino acid and if there were any differences in carbohydrate preference among clinical UPEC. Thus, we analyzed growth in LB, pooled human urine, and M9 minimal medium with five different sole carbon sources including casamino acids (CAA), glucose, glycerol, galactose, or ribose (**Fig. S1A-G**). These individual data were used to calculate the area under the curve (AUC) to display differences more easily between strains (**Fig. 1C**).

Unsurprisingly, we found robust growth in LB for all *E. coli* clinical isolates as well as CFT073 and K12 (**Fig. 1C)**. However, we observed a remarkable defect in K12 versus all UPEC isolates when grown in human urine (**Fig. S1G**); potentially growth in urine could differentiate between *E. coli* strains that can infect the urinary tract. Interestingly strains HM17, HM43, and HM68 had restricted growth on all sugar carbon sources (**Fig. 1C**). However, these strains had robust growth when given CAA as a sole carbon source, as did the rest of the UPEC strains (**Fig. 1C**). This result further corroborates the studies that show UPEC prefers amino acids as the optimal carbon source *in vivo* (31, 35).

Iron, a vital nutrient required for bacterial growth, is highly restricted within the host (36, 37). Most UPEC strains encode several different iron acquisition systems that include the synthesis of siderophores to acquire iron sequestered by the host (29, 38). We assessed siderophore production of these clinical isolates using the Chrome Azurol S (CAS) assay (39). Strains were cultured overnight in iron-poor M9 medium, then their supernatants were incubated with CAS dye, which is conjugated with Fe^3+^. When Fe^3+^ is chelated from the dye, it changes from blue to red, resulting in a loss of absorbance at 630 nm. We observed variable levels of CAS activity between strains. Strains CFT073, HM6, and HM14 had strong CAS activity, chelating the dye at levels similar to the positive control EDTA (**Fig. 1D**). Six of the strains, HM54, HM56, HM57, HM66, HM68, and HM86, displayed more modest CAS activity (**Fig. 1D**), though still significantly different from the medium-alone control. The remaining five UPEC isolates, strains HM3, HM7, HM17, HM43, and HM68, as well as K12, had no significant CAS activity (**Fig. 1D**). Apart from *E. coli* K12, it was surprising to observe this apparent lack of activity from strains that are pathogenic and all encode siderophore systems. Therefore, we adjusted the assay and cultured the strains directly on agar plates containing CAS. While the CAS agar does not result in quantitative data, it is more sensitive to siderophore production since signal builds as the colonies grow and secrete siderophore (39). Using CAS agar, we detected siderophore production in all strains, including K12 (**Fig. S1H**).

### Surface structure composition varies between isolates, resulting in unique motility, biofilm, and morphology phenotypes

Uropathogens use motility to ascend the ureters from the bladder to the kidneys and to move to nutrient-rich areas (40-42). To assess the motility potential of our clinical collection of UPEC strains, we conducted a motility assay in semi-soft agar at 30°C (**Fig. S2A**). After 16 hours of incubation, the swimming diameter was measured (**Fig. 2A**). Interestingly, four strains were non-motile: HM14, HM54, HM57 and HM86 (**Fig. 2A**). On the other hand, HM3, HM6, HM7, HM43, HM56, and HM66 were hypermotile compared to both CFT073 and K12 (**Fig. 2A**). While the ability to move between organ sites is important, so is the ability to establish a permanent community. Uropathogens do this through cell-to-cell interactions, both on abiotic structures such as catheters and within eukaryotic cells as intracellular bacterial communities (IBCs) (43-45). We tested biofilm formation in the clinical isolates at both 30°C and 37°C using a microtiter plate assay (**Fig. 2B**). In LB at 37°C, only HM6, HM17, HM57, and HM68 produced biofilm similar to CFT073 (**Fig. 2B**). It is notable that at 37°C, we observed the biggest differences between CFT073 and K12 as compared to 30°C. At 30°C, HM3, HM43, HM66 and CFT073 produced the largest biofilm communities (**Fig. 2B**). Urine biofilms were barely above background detectable levels (**Fig. S2B**). In addition to examining biofilm potential via crystal violet staining, we also assessed curli formation through growth on a medium containing Congo red dye (**Fig. 2C**). This dye binds curli, which aid in biofilm formation: curli-positive bacteria appear red and curli-negative colonies appear white (46). HM14 displayed a unique morphology that very clearly excluded the Congo red dye and exhibited a mucoid phenotype (**Fig. 2C**). HM3, HM7, HM56, HM68, and K12 all displayed a very dark red color (**Fig. 2C**). Many of the strains displayed moderately red or pink colors, indicating a spectrum of curli presence among uropathogens under these conditions.

**FIG 2.**
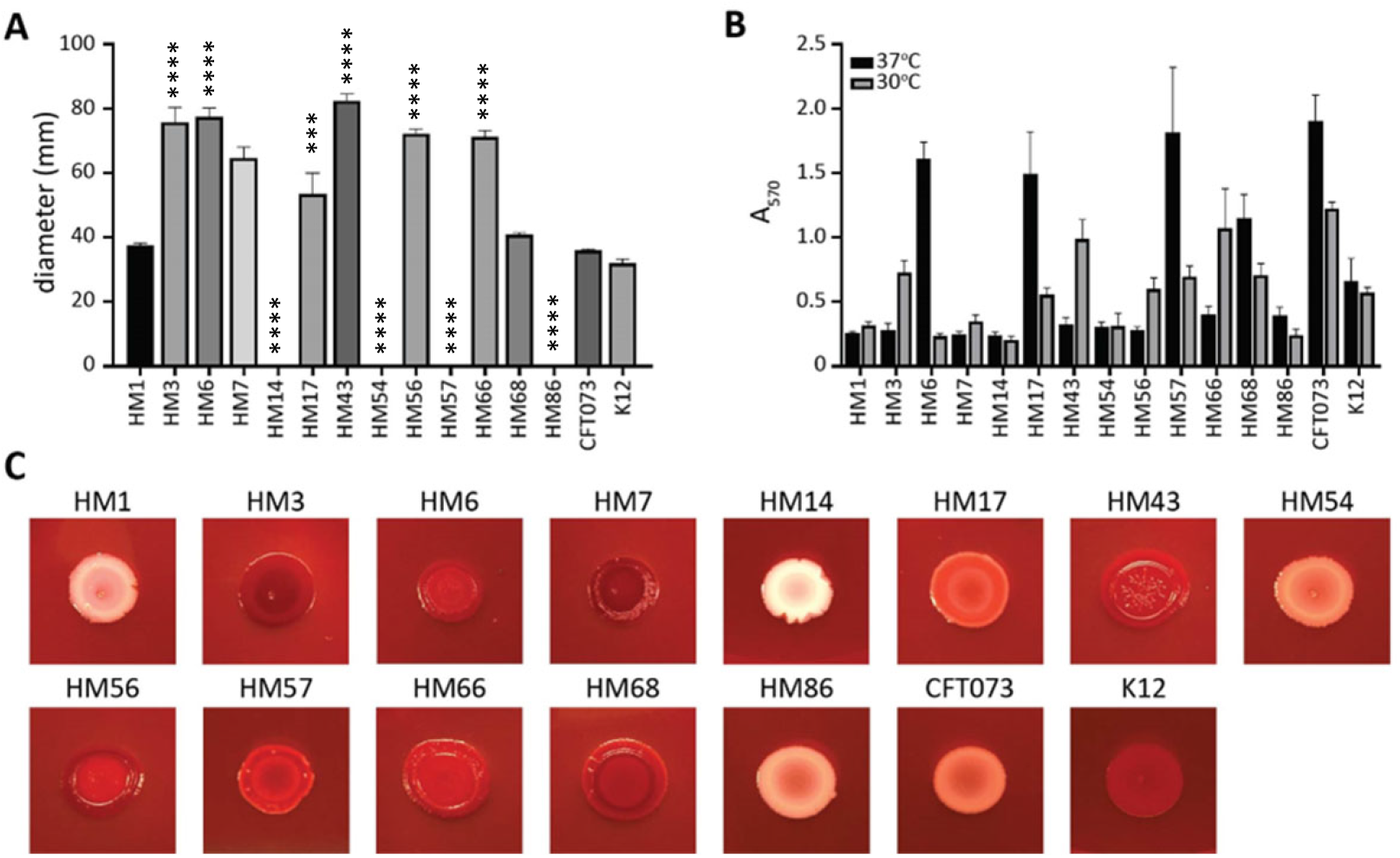
Some UPEC isolates are unable to swim or form biofilms *in vitro*. (A) Motility was assessed by stabbing clinical isolates into semi-solid agar and incubating at 30°C for 16 h. Measured diameters are shown in mm with error bars to indicate SEM. Bars represent the average of biological replicates (n=3). One-way ANOVA was performed with Dunnet’s multiple test correction compared to CFT073. ***, *P* < 0.005; ****, *P* < 0.0001. (B) The ability of clinical UPEC isolates to form biofilms in LB was determined at both 37°C (black) and 30°C (grey). Bar height represents the average absorbance (570nm) after staining with crystal violet of four biological replicates, error is SEM. (C) Overnight liquid cultures were spotted onto LB agar plates containing Congo red dye and incubated for 48 h at 30°C.

### Type 1 fimbrial expression varies greatly among clinical isolates

Adherence is a critical factor during UTI. Specifically, the use of type 1 fimbriae to bind the uroepithelium is essential to withstand the flow of urine (47). These fimbriae are phase-variable via invertible element (48). We utilized a PCR-based assay to determine whether UPEC strains had their invertible elements in the ON or OFF orientation (49) and assessed the percentage of the total bacterial population with the invertible element in the ON position after being cultured statically at 37°C, a condition that has previously been shown to induce ON switching (49). HM17 and HM56 had over 50% of the invertible element in the ON position, while HM6, HM14, HM54, HM57, and HM86 had nearly undetectable *fim*-ON populations (**Fig 3A**).

**FIG 3.**
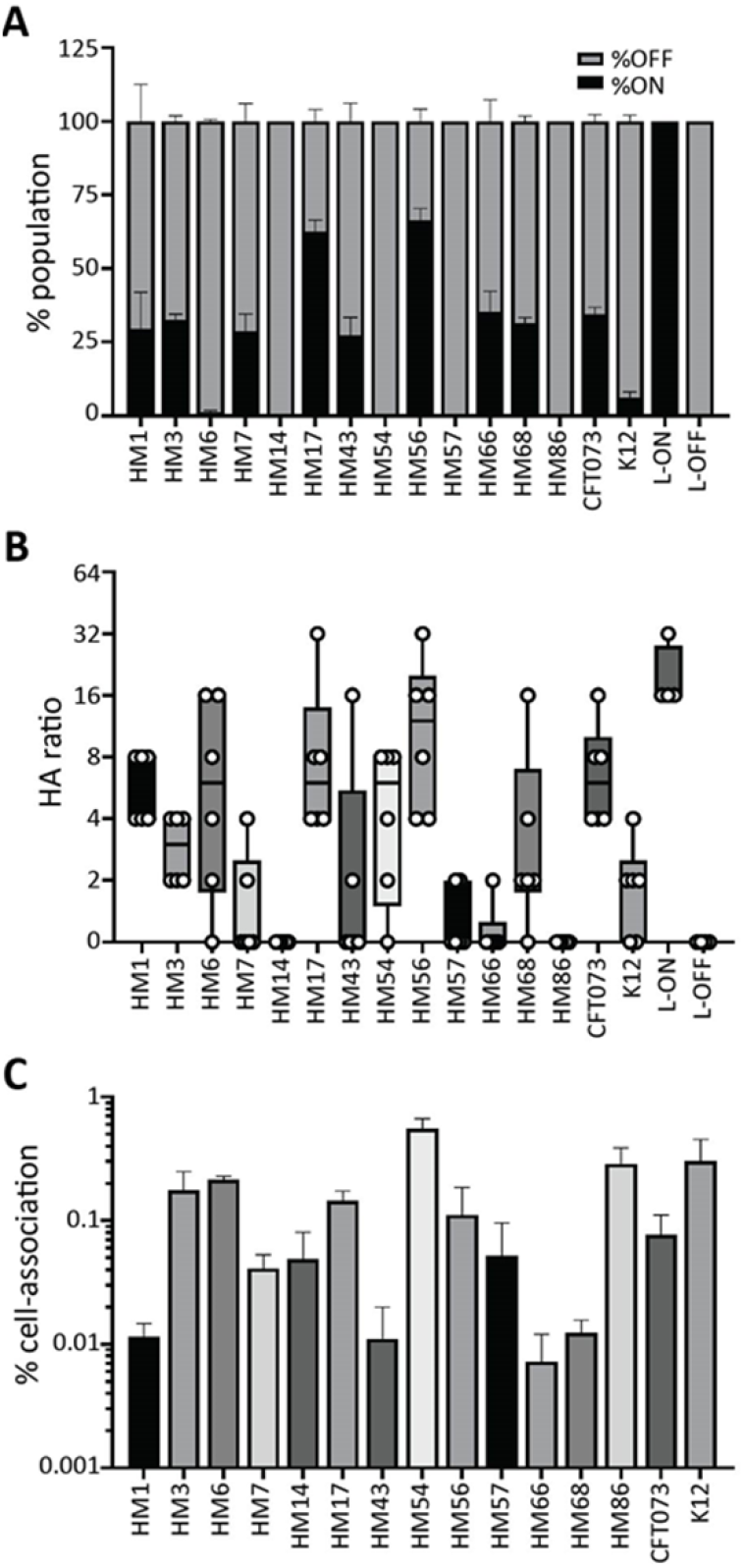
HM17 and HM56 demonstrate increased type 1 fimbrial expression concordant with hemagglutination and adherence phenotypes. (A) The invertible element (IE) PCR assay was used to quantify the orientation of the *fimS* IE as either on or off. The stacked bar height indicates the mean of the population with *fim* ON (black) and *fim* OFF (grey). Data represent three biological replicates with SEM shown. CFT073 L-ON and L-OFF constructs were used as controls in which 100% of the total population should be either *fim* ON or OFF. (B) Hemagglutination (HA) assays were performed with guinea pig erythrocytes utilizing bacterial titers in a 2-fold serial dilution series, indicated by the HA ratio. Box and whisker plots denote the quartiles and ranges of the data from six biological replicates, shown by the individual points. CFT073 L-ON and L-OFF strains were used as controls. L-ON can hemagglutinate at high ratios (1:32), indicating that fewer bacteria are required to produce the phenotype. (C) Clinical UPEC strains were assessed for their ability to adhere to the T24 bladder cell line. Cell-associated bacteria were enumerated after 1 h of co-culture with cell monolayers; the bars indicate the mean of four biological replicates with SEM error bars. Data shown represent the normalized output to input ratios as a percentage.

An alternate method to determine the expression and functionality of bacterial fimbriae is through a hemagglutination assay (14). This quantifies UPEC adherence to guinea pig erythrocytes. Bacteria were serially diluted before adding erythrocytes; a higher ratio indicates stronger type 1 fimbrial expression. HM1, HM6, HM17, HM56 and CFT073 all exhibited strong hemagglutination phenotypes (**Fig. 3B**). HM14 and HM86 had no ability to agglutinate red blood cells under these assay conditions (**Fig. 3B**). This phenotype is type 1 fimbria specific, and the addition of mannose was sufficient to inhibit all hemagglutination (**Fig. S3B**).

Besides type 1 fimbriae, UPEC employ an arsenal of other adhesins to bind epithelial cells (50). To quantify the contribution of all these systems we added bacteria to bladder cells at a multiplicity of infection (MOI) of 100 and incubated for 1 hour before determining the number of cell-associated CFU. The ability of clinical isolates to associate with cells widely ranged from 0.09% (HM66) to 0.75% (HM54) when compared to input CFU (**Fig. 3C**). Interestingly, non-pathogenic K12 had higher cell-association than CFT073 (**Fig. 3C**). Also of note, while HM14 does not appear to have the type 1 pili (**Fig. 3A, B**) it was able to successfully cell-associate *in vitro* (**Fig. 3C**). This demonstrates that there are other, likely surface-expressed structures, that aid in UPEC-cell association (50).

### Host cell death is directly linked to *alpha-*hemolysin

The ability of UPEC to kill host cells is initiated by a variety of known virulence factors. For example, hemolysin and other secreted toxins, are known to disrupt host cell viability and cause dysregulation to essential host cell signaling cascades (51, 52). To investigate the effect of clinical UPEC on host cell viability, we measured their interactions with bladder and kidney cell lines (**Fig. 4A, B**). We used a colorimetric MTT assay which assesses the viable cell number. Monolayers of host cells were treated with an MOI of 50 for 5 h before determining host cell viability. CFT073 and HM86 were the only strains that caused cell death in T24 bladder cells (**Fig. 4A**): Both strains were the only two that carry *alpha*-hemolysin. HM43 and 66 seemed to have increased cell viability, but upon further investigation, we observed larger biofilm-like bacterial populations that appeared to be viable even after antibiotic treatment. Since the MTT can be reduced by any living organism, we had unexpected background in our assay. We observed that HM86, CFT073 and HM56 were able to kill HK2 kidney cells in culture (**Fig. 4B**). It was interesting that the only two strains carrying *alpha*-hemolysin were able to kill both cell lines, while the *beta*-hemolytic HM56 was only able to kill kidney cells. These results were supported by our findings on blood agar, where only HM56, HM86, and CFT073 were able to lyse red blood cells (**Fig. 4C**). Other UPEC-associated toxins such as Vat, Sat, Pic, Tsh, and CNF1 were present only in a few strains and their presence did not correlate with cell viability.

**FIG 4.**
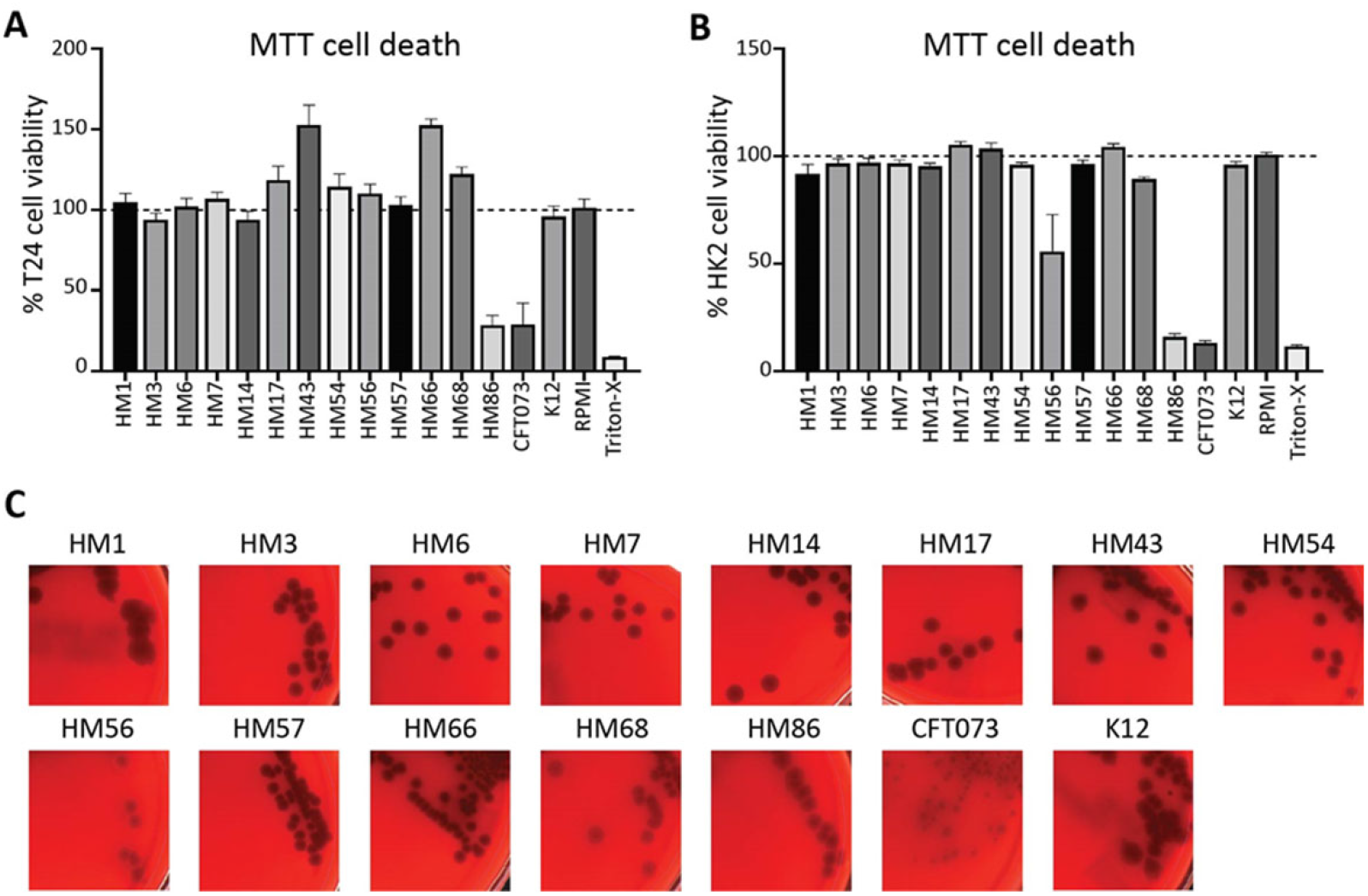
HM56 and HM86 uniquely carry *alpha*-hemolysin and can kill uroepithelial cells in culture. To determine the ability of clinical UPEC strains to lyse epithelial cells, monolayers of (A) T24 bladder or (B) HK2 kidney cell lines were treated with UPEC for 5 h. The MTT cell viability kit was used to measure monolayer viability pre- and post-treatment with UPEC. Cellular respiration causes a color change, which can be measured via absorbance. These data were normalized and mean values of four biological replicates are indicated by the bars with SEM. RPMI cell culture medium with no bacteria served as a negative control and 0.4% Triton-X 100 served as a positive control for cell death. (C) UPEC strains were struck onto blood agar plates and incubated overnight at 37°C. The lysis of red blood cells leads to a zone of clearance around isolated bacterial colonies. CFT073 was used as a positive control for hemolysis.

### Assessment of stress response phenotypes reveals diverse strain responses

Neutrophil bombardment is one of the host’s first lines of innate defense against UTI (53). Neutrophils attack bacteria by releasing reactive oxygen species (ROS) contained in their granules. Survival in H_2_O_2_ is designed to assess the viability of bacteria in the presence of extracellular ROS. A kill curve for each strain was determined by plotting the mean CFU/mL every 15 minutes for 1 hour in 0.2% H_2_O_2_ (**Fig. 5A**). Three strains (HM3, HM7 and K12) were not killed by H_2_O_2_ (**Fig. 5A**). HM68 was the most highly affected strain, with a 10,000-fold loss of CFU over sixty minutes (**Fig. 5A**). Increased susceptibility to death via ROS could suggest a weaker *in vivo* survival rate during infection; however, shockingly, commensal isolate K12 was one of the most resistant strains (8-fold reduction in CFU), indicating that this phenotype may not be predictive or all-encompassing of the immune interaction experienced during UTI.

**FIG 5.**
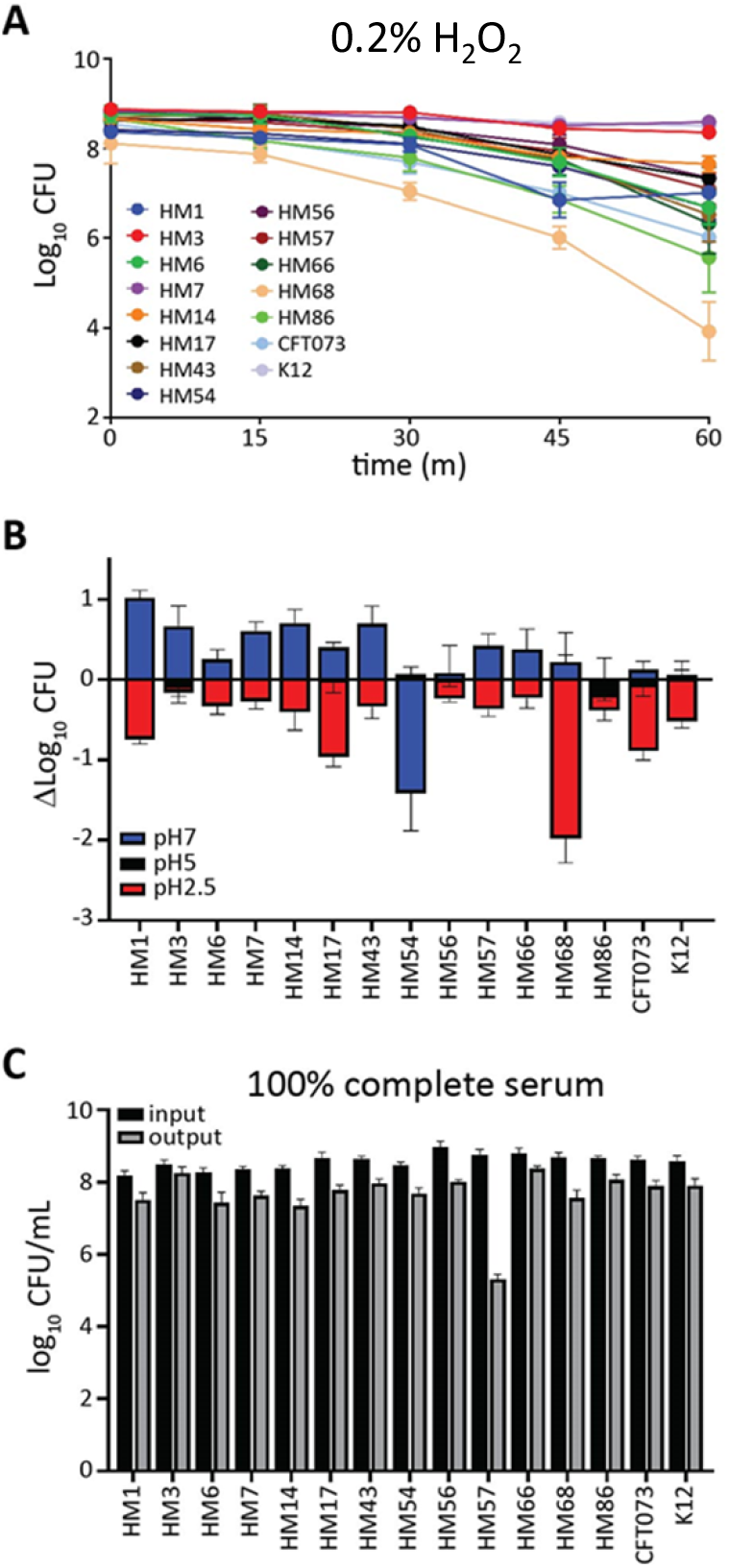
UPEC isolates are mostly resistant to peroxide, acid, and serum stresses HM68 has decreased resistance to hydrogen peroxide and acid stress, while HM57 only displays deficiency in human serum resistance. (A) 10^8^ CFU/mL of UPEC were inoculated into LB containing 0.2% H_2_O_2_ and maintained statically at room temperature. Samples were taken and serial diluted at 0, 15, 30, 45, and 60 min to enumerate CFU/mL. The means of four biological replicates with associated SEM are shown. (B) From an overnight culture, UPEC were diluted to 10^8^ CFU/mL in LB buffered with MES to pH 7, 5, or 2.5. Cultures were incubated for 2 h at 37°C with aeration and CFU/mL were quantified at both 0 and 2 h to calculate the change in CFU/mL. Error bars represent SEM calculated from the mean of four biological replicates. (C) UPEC cultured overnight in LB and washed and resuspended in PBS, then 10^8^ CFU/mL was added to 100% complete human serum. Samples were incubated statically at 37°C for 90 minutes, and both input (black) and output (grey) samples were enumerated and plotted. Bars represent the mean of four biological replicates with SEM error bars.

When in the urinary tract, bacteria are exposed to a variety of host-mediated stress responses including osmotic stress, pH changes, bombardment from the immune system, and nutrient limitation. It is plausible that pathogens may have enhanced responses to these environmental stressors. Therefore, we performed a series of assays to investigate UPEC stress responses. First, we examined UPEC growth and survival after a one-hour exposure to LB buffered to pH 7, 5, and 2.5 (**Fig. 5B**). We observed that most strains had growth up to 10-fold or had no change in CFU in pH 7. The exception was HM54, which experienced over a 14-fold decline in population (**Fig. 5B**). Interestingly, this strain was one of the most tolerant to pH 2.5, along with HM3 (**Fig. 5B**). Most strains experienced a 10-fold or less reduction in CFU/mL in acidic media except for HM68 which had a 20-fold loss (**Fig.5B**). Under these conditions, *E. coli* K12 was not exceptionally different from the average of UPEC strains.

Complement-mediated killing is also an important immune defense that can be tested by measuring bacterial survival in human serum. We treated the strains with 100% human serum for 90 minutes and enumerated CFU. Only a single strain, HM57, was susceptible to serum (**Fig. 5C**) with a loss of about 10^4^ CFU compared to input. This loss of CFU was not observed when HM57 was incubated with heat-inactivated serum (**Fig. S4**), indicating the killing was complement-mediated. K12 was comparably resistant to serum, a surprising result, given that it is a commensal isolate.

### Murine model of ascending UTI was used to assess strain infectivity

While these strains were isolated from women with symptomatic UTI making them pathogenic (30, 31), we had not yet tested their ability to establish disease in our mouse model of UTI. We infected animals with each of the clinical isolates as well as CFT073 and K12 as well as using colonization data of HM43, 56 and 86 from a previous study. At 48 hours post inoculation, we enumerated bacterial burden in the urine, bladder, kidneys, and spleen (**Fig. 6, S5**). Each strain was able to colonize at least one site, and most strains were able to colonize to similar levels as CFT073 (**Fig. 6**). However, some strains displayed a preference for specific organ sites. Overall, the spleen was the least colonized organ site in which only seven strains were able to achieve detectable CFU (**Fig. S5**). We were unable to recover CFU from the bladders of mice infected with HM14, but both the urine and kidneys of these mice were highly colonized (**Fig. 6**). This observation is especially interesting given that HM14 does not encode type 1 fimbriae, which mediate binding to bladder epithelium (54). In contrast, HM66 robustly colonized the urine and bladders of mice, but was absent in the kidneys. The differences in colonization between these strains, and specifically the preference for particular organs could be illuminating in dissecting the virulence factors essential to these tissue-specific advantages.

**FIG 6.**
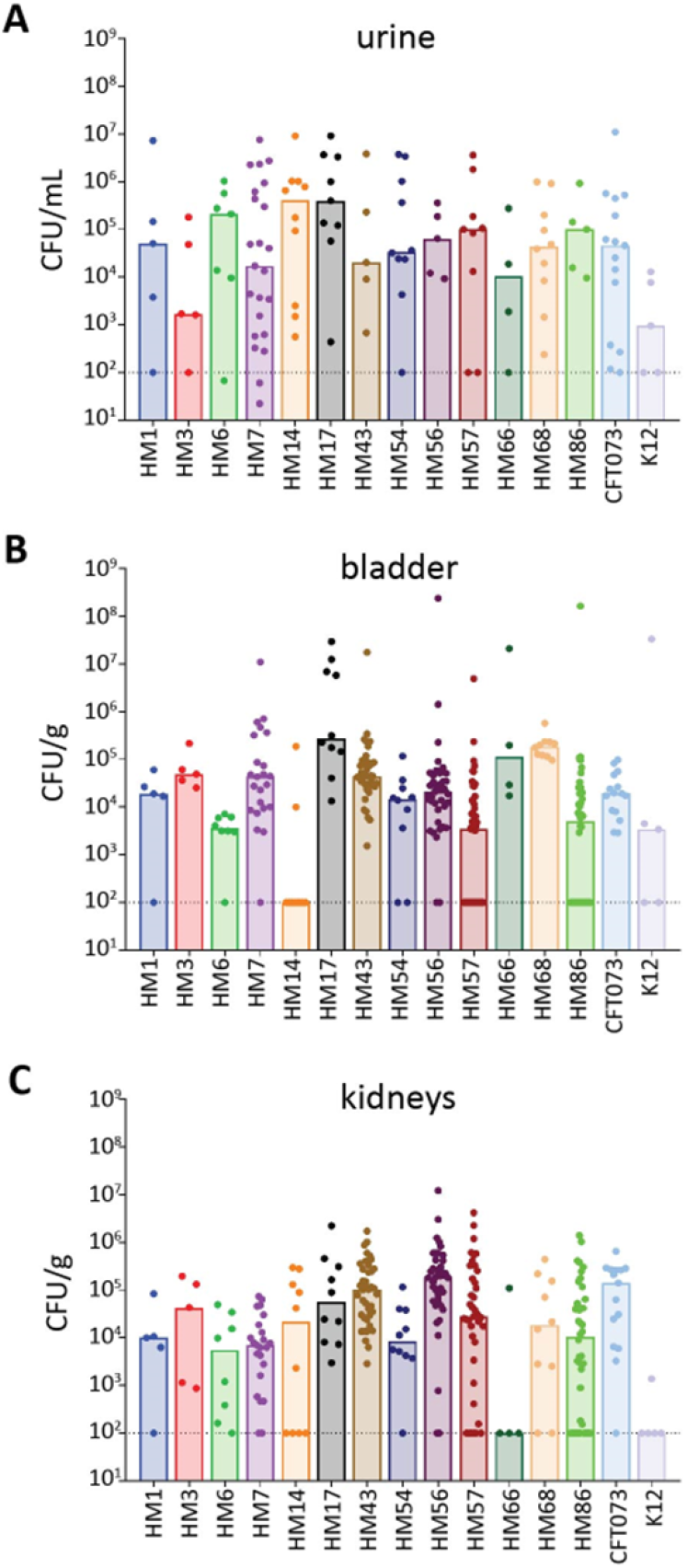
Human UPEC isolates readily establish UTI in mice. 6–8-week-old CBA/J mice were transurethrally inoculated with 10^8^ CFU of UPEC. The infection progressed for 48 hrs before urine, bladder, and kidneys were harvested to quantify bacterial burden in each organ site. The limit of detection is 100 CFU per g of tissue or mL urine and is indicated by the dashed line; bars denote the median of bacterial burden. Each strain was tested on 5-40 CBA/J mice, dots represent individual mice. HM43, HM56, and HM86 were previously published (31, 34). (A) Urine samples were collected from each mouse and brought up to a final volume of 150µL in sterile PBS. CFU/mL of urine was determined for each individual mouse. (B) Bladder and (C) kidneys were aseptically harvested and homogenized in 3mL of sterile PBS, then plated on LB agar to determine CFU/g of tissue.

### Mathematical modeling of bacterial burden using phenotypic data to develop predictors of UPEC and UTI severity

Finally, we wanted to create a holistic model to predict infectivity using this wealth of phenotypic data. We used multiple linear regression to predict bacterial burden in the mouse model, which is acting as the proxy for infectivity.

However, this model is only predictive of normally distributed data. Therefore, using the Shapiro-Wilk test, we determined if the urine, bladder, and kidney colonization data were normally disturbed. We found that only the bladder and urine data were (**Fig. S6A**), so, unfortunately, we could not use this model to predict kidney colonization. Another stipulation when using multiple linear regression is that none of the input factors can be correlated with one another. Therefore, we created a correlation matrix between all 18 phenotypic assays we performed (**Fig. S6B**), with a cutoff r^2^ value of 0.5 as “correlated.” We found that all growth conditions in M9 correlated with one another, as did acid sensitivity. Growth in urine correlated with all M9 growth conditions, except when glucose was the sole carbon source. We also observed expected correlations; % *fim* ON correlated with both hemagglutination and motility.

From there, we correlated each of these phenotypic assays with either urine or bladder colonization (**Fig. 7A, Suppl. Table 1**). Some factors were clearly, strongly correlated with colonization. For example, type 1 fimbrial expression was highly correlated with bladder colonization, and growth in *ex vivo* urine correlated with bacterial burden in urine during experimental UTI. Guided by these single regressions, as well as manual curation based on previous studies, we used type 1 fimbrial expression, siderophore production, and growth in M9 with CAA as the variables for bladder colonization (**Fig. 7B**). The adjusted R^2^ for the multiple linear regression model was 0.6411, compared to the single R^2^ values of 0.4913, 0.4319, and 0.3075 for used type 1 fimbria expression, siderophore production and growth in M9 with CAA, respectively (**Fig. 7C**). For urine colonization we used growth in *ex vivo* urine, hemagglutination, and motility (**Fig. 7B**). The adjusted R^2^ for this combined model is 0.4821, a robust increase from 0.2728, 0.1864, and 0.08827 for *ex vivo* urine, hemagglutination, and motility individually (**Fig. 7C**). Altogether, this underscores the wide diversity of virulence strategies that UPEC can utilize, and how this diversity might be driving a corresponding difference in tissue tropism and fitness.

**FIG 7.**
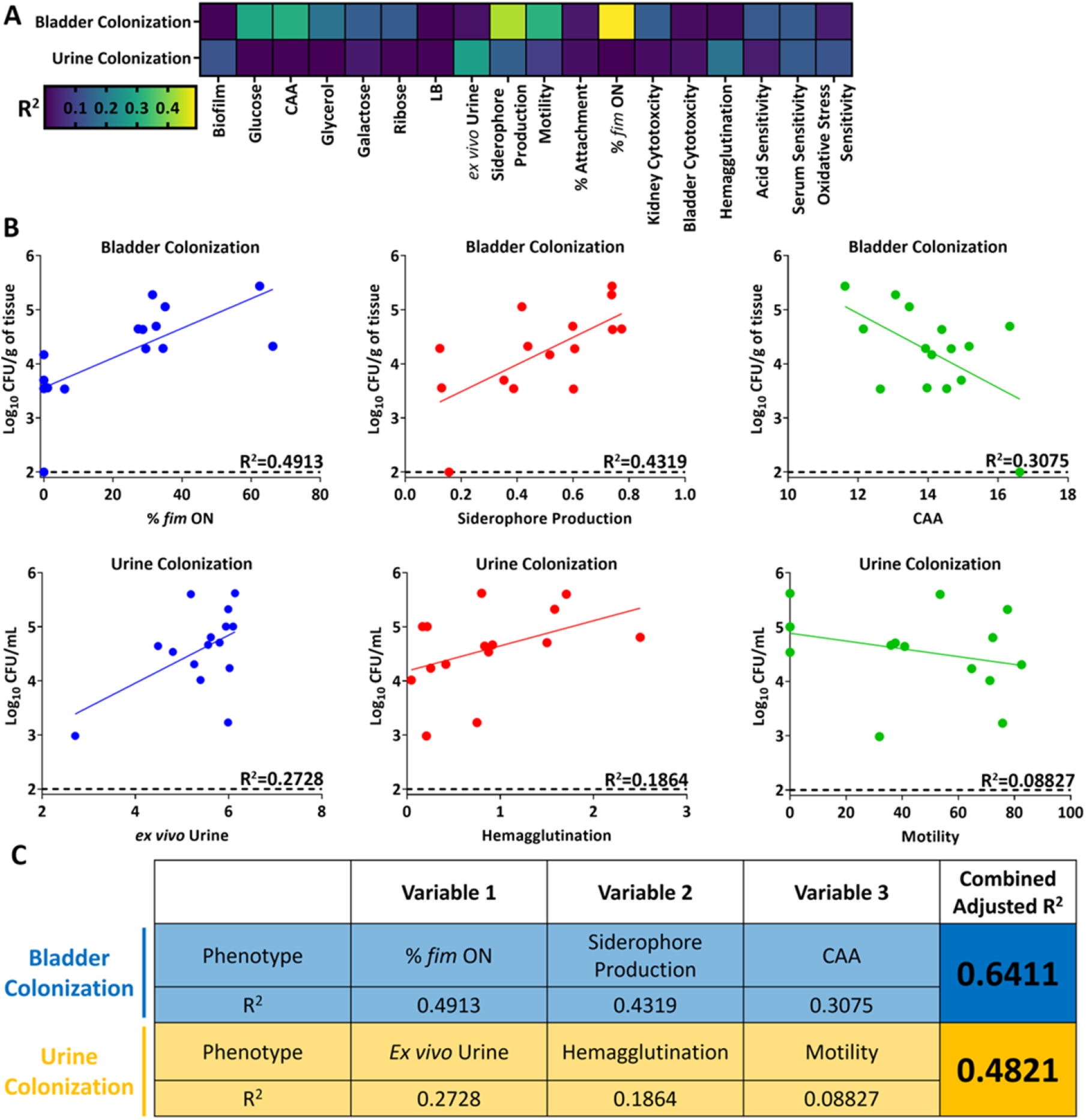
Multiple linear regression can partially predict levels of bladder and urine colonization. (A) Pearson’s correlation between either bladder colonization or urine colonization for each indicated phenotype assay. (B) Select phenotypes used in multiple linear regression with displayed R^2^ values based on Pearson’s correlation. (C) Multiple linear regression used to model either bladder or urine colonization. Each variable used in the model is displayed with its corresponding single R^2^ value, as well as the final, adjusted R^2^ value when all three variables are taken into account.

## Discussion

UPEC are an incredibly diverse group of pathogens, among an already extremely varied species (55, 56). This population causes a variety of disease pathologies, including uncomplicated, complicated (*e*.*g*., catheter-associated), asymptomatic, and recurrent urinary tract infections. Accordingly, UPEC isolates encode a large accessory genome where most uropathogen-specific genes exist. Interestingly, we have found that some of our clinical isolates do not even possess the well-established genes that are generally considered to be essential for UPEC pathogenesis, such as type 1 fimbriae (38), though the redundancy of the UPEC genome could be compensating for the lack of these virulence factors. Alternatively, these clinical isolates lacking traditional virulence factors could be encoding novel proteins absent within UPEC prototype strains. In order to understand the virulence strategies of these uncharacterized strains, we decided to take a functional approach, rather than genetic since essential virulence mechanisms often have corresponding phenotypes. We hypothesized that using phenotypic assays as opposed to genotyping would be more revealing of the uropathogenic potential of these strains.

Therefore, we performed an extended series of phenotypic microbiological assays to quantitatively describe bacterial urovirulence traits common to UPEC. We then used linear regression to correlate phenotype outcomes to bacterial burden in an ascending model of murine UTI to quantify disease severity. We found growth in minimal medium with CAA as a sole carbon source, siderophore production, and type 1 fimbrial expression were able to best predict bladder colonization outcomes. Urine outcomes were best modeled by bacterial motility, hemagglutination, and growth in human urine *ex vivo*. Unfortunately, although we determined the bacterial burden in the kidneys and spleen (indicating bacteremia), we were unable to perform this correlation analysis since the data were not normally distributed.

Our data suggest that growth in minimal medium with CAA as a sole carbon source is a good indicator of bladder colonization. Interestingly, this correlation was negative even though UPEC prefer amino acids as carbon sources during infection (35, 57). However, CAA are a mix of single amino acids and nutrient milieu in the infected bladder is far more likely to contain oligopeptides or even whole proteins from lysed cells. Indeed, several di- or tri-peptide transporters are virulence factors in the mouse model (35). To use CAA as a carbon source, UPEC would need to rely on the upregulation of single amino acid transporters, instead of less specific di- or tri-peptide transporters as observed *in vivo*.

We also observed three clinical strains, HM17, 43, and 68, had restricted growth with sole carbohydrate sources in minimal medium. However, these strains had some of the highest bacterial burdens in the bladders of mice, showing that that sugar utilization is dispensable for UPEC in the infection niche, corroborating several previous studies (31, 35, 57). It is interesting to speculate that these strains might have defects in gut colonization where sugar utilization is key (58).

Iron sequestration mechanisms are highly expressed during infection (30, 31, 59), but are not part of the core genome of *E. coli* (31). There are four main UPEC siderophore systems used for scavenging iron in the host: yersiniabactin, aerobactin, enterobactin and salmochelin (27, 29, 60). The clinical strains encoded between one to three of these, while CFT073 encodes three. We observed differences between strains in liquid CAS assay, but every strain had CAS activity when assessed by the more sensitive agar-based method. Interestingly, strains with lower liquid CAS activity had higher bladder colonization (HM17, HM43 and HM68). Siderophore production comes at a high metabolic cost, and perhaps these lower producing strains have fine-tuned production of these small molecules while the others are over-producing.

Type 1 fimbriae are a preeminent virulence factor for murine bladder adhesion, invasion, and colonization (61). An invertible element assay determined the orientation of the type 1 fimbrial promoter in either the on or off orientation *in vitro*, and the results were highly predictive of bladder colonization in the murine model. Correspondingly, we were unable to recover any CFU from the bladders of mice inoculated with HM14, which does not encode type 1 fimbriae. However, this strain was able to robustly colonize the urine and the kidneys of the mouse. Thus, although type 1 fimbriae are critical for establishing cystitis, they are dispensable for colonization of other sites in the urinary tract. The presence of the *fim* operon is necessary but not sufficient to predict bladder colonization, as non-pathogenic K12 also possess this fimbria. However, by using the invertible element assay, we revealed that the invertible element was overwhelmingly in the off position in K12. This would phenocopy a strain that does not encode the fimbriae, demonstrating the utility of phenotypic testing over genotyping.

HM14 was particularly unique in more ways than its lack of the *fim* operon. It also had a more mucoid appearance, complete exclusion of Congo red dye, and a lack of motility. The lack of motility is particularly interesting given the strain’s high kidney colonization considering previous literature demonstrates the critical role of flagella-mediated motility in ascension of the urinary tract (40). However, we must consider *in vitro* conditions such as temperature, nutrient availability, and presence of ROS, among other factors, are different compared to the murine host. This may lead to different expression levels of flagella in the host environment.

Overall, we found the single phenotype correlations relating back to urine colonization were weaker than the ones for bladder colonization. The strongest single correlation was a positive correlation with human urine growth *ex vivo*, a logical observation given the infection milieu of the urinary tract is urine. However, it is worth noting that the data are skewed; K12 has a severe defect in growth and colonization in urine, while the UPEC strains are clustered more tightly together (**Fig. 7B**).

Interestingly, bacterial motility had a mild negative correlation with urine colonization. This could indicate that more motile strains are better able to move and associate with the bladder epithelium, ascend into the kidneys, or are more highly recognized by the TLR5-mediated host clearance mechanisms; any or all of these are possible.

One of the relatively stronger associations was a positive correlation between bacterial hemagglutination of guinea pig red blood cells and urine colonization. Likewise, a previous study found hemagglutination to predict bladder colonization in C3H/HeN mice (8). They also speculated that some of the observed differences between *in vivo* and *in vitro* conditions were likely due to differential responses to host niche-specific environmental cues (8). It is known that bladder epithelial cells are shed into the urine as a host defense mechanism to infection, so it is plausible that host cell-bound bacteria could be concentrated in the murine urine sample, also explaining the association.

Collectively, we demonstrate that prediction of virulence potential and assessment of infection severity is better determined through phenotypic analyses. The current limitations in the potential clinical implementation of these assays come from expense, time, and a lack of standardized experimental conditions that more accurately model the host. However, the use of these current phenotypic characterization assays increases the prediction efficiency of UPEC pathogenic potential. Future studies will work to improve these assays and make them more reflective of the host environment, hopefully resulting in even better predictive outcomes.

## Materials and Methods

### Bacterial and cell culture conditions

*E. coli* CFT073 was isolated from the blood and urine of a hospitalized patient with acute pyelonephritis (20). Lysogeny broth (LB), which contains 0.5 g NaCl, 5 g yeast extract, and 10 g tryptone/L, was used to routinely culture bacteria at 37°C under aerated or static conditions and was inoculated from single colonies.

Human epithelial T24 bladder cells (ATCC HTB-4) and HK2 kidney cell (ATCC CRL-2190) were maintained in RPMI medium containing L-glutamine (Corning 10-040-CV) with 10% fetal bovine serum (FBS) and (10mg/mL) penicillin/streptomycin antibiotics at 37°C and 5% CO_2_.

### Bacterial growth curves

Overnight LB bacterial cultures were washed once with phosphate-buffered saline (PBS), then inoculated 1:100 into 1 mL LB, or M9 minimal medium (62) supplemented with either 0.4% glucose, glycerol, galactose, ribose or casamino acids (CAA), or into filter-sterilized pooled human urine collected from at least four female donors. These cultures were incubated with aeration at 37°C in a Bioscreen-C automated growth curve analyzer (Growth Curves USA), collecting OD_600_ readings every 15 min for 24 h. Area under the curve (AUC) was calculated using GraphPad 9.3.1.

### Anaerobic sugar fermentation

The protocol was adapted from Himpsl *et al*. 2020 (63). Briefly, 5 µL of overnight LB cultures of each strain was spotted onto LB agar containing phenol red (0.04 g/liter) alone, or supplemented with 0.4% glucose, 0.4% glycerol, 0.4% galactose, or 0.4% ribose. Plates were incubated for 24 hours at 37°C under anaerobic conditions (BD GasPak EZ Anaerobe) and then imaged. Plates were transitioned to aerobic conditions and incubated at room temperature for another 6 or 24 hours, imaging at each time point. In parallel, strains were also spotted on plates that were incubated in aerobic conditions at 37°C for 24 hours, before transitioning to room temperature for another 6 and 24 hours. Colors were quantified from images of representative plates. The center of each colony was selected with the eyedropper tool in Adobe Photoshop to determine its color.

### Chrome Azurol S assay

Strains were cultured overnight in LB with shaking at 37°C. The overnight cultures were harvested by centrifugation, washed with PBS, and diluted 1:100 into M9 minimal medium supplemented with 0.4% glucose and incubated overnight, shaking at 37°C. Cultures were pelleted by centrifugation, and 100 µL of supernatants were combined with 100 µL of chrome azurol S (CAS) shuttle solution (64) and incubated at room temperature for 30 minutes. 10 mM EDTA served as a positive control. After 30 minutes, the absorbance at 630 nm was recorded.

### Biofilm formation

Strains were cultured overnight in LB, shaking at 37°C. and diluted 1:100 into either 3 mL of fresh LB medium, or *ex vivo* human urine into tissue culture treated 24 well plates (Corning), and incubated statically at either 37°C or 30°C for 24 hours. The medium was carefully decanted and rinsed with distilled water to remove any non-adherent bacteria, then stained with a 0.1% solution of crystal violet for 15 minutes at room temperature. The stain was aspirated and wells rinsed with distilled water. Biofilms were solubilized with 1 mL of 100% ethanol for 15 minutes at room temperature with shaking. 200 µL of solubilized biofilm was read in a 96-well plate at an absorbance of 570 nM. Congo red agar was made as previously described (65). Overnight cultures (5μL) were spotted onto Congo red agar and incubated at 30°C for 48 h before images were taken.

### Motility

Overnight LB cultures of each strain were normalized to an OD_600_ of 1.0 and resuspended in HEPES buffer, pH 8.4. Cultures were stabbed in duplicate into tryptone agar plates (1% tryptone, 0.5% NaCl) containing 0.25% agar with an inoculating needle. Bacteria were allowed to swim for 16 hours at 30°C and then the diameter of the motility zone was measured.

### *fimS* invertible element orientation

Bacteria were cultured overnight in LB medium statically at 37°C. The OD_600_ of cultures was taken and strains were normalized to OD_600_ 0.5 in water and PCR, product digestion was done as previously described (49). The entire sample was run on a 3% agarose gel at 100 V to visualized *fimS* ON and OFF orientation bands. A representative gel is shown in **Fig**. S3A. The intensity of the bands was quantified using Biorad software.

### Hemagglutination

Bacteria were cultured statically in LB at 37°C for 72 hours (53). Cells were washed and resuspended in 1X PBS then serially diluted 1:2 in a round-bottom 96-well plate. Guinea pig erythrocytes were washed, resuspended at 3% in PBS (vol/vol), then added to each well of the plate. Bacteria and erythrocytes were gently mixed and allowed to settle for one hour. To assess the role of the type 1 fimbriae, 50 mM mannose was added to interfere with fimbriae binding as previously described (57).

### Cell association

Cell lines were cultured to confluence in 24-well plates (Corning) and serum-starved overnight before performing adhesion assays. Bacteria were cultured overnight in static LB at 37°C. Cells were trypsinized and counted using a hemacytometer, and bacterial CFU/mL of overnight cultures was determined via OD_600_. A multiplicity of infection (MOI) of 1:100 was used for these assays with bacteria normalized in 1mL serum-free RMPI medium and added to epithelial cell monolayers for a 1 h incubation. Bacteria-containing medium was aspirated, and cell monolayers were gently washed with 1X PBS three times with mild agitation. A PBS solution with 0.4% Triton-X 100 was applied for 30 minutes with strong agitation to lyse epithelial cells. Resulting lysates were serially diluted and plated on LB agar to enumerate CFU/mL of cell-associated UPEC.

### MTT cell survival

Cell viability after co-incubation with *E. coli* was determined utilizing the Cell Proliferation Kit I (Milipore Sigma #11465007001). A suspension containing a MOI 50 of each strain was added to a confluent monolayer of host cells (T24 ATCC or HK2 ATCC) and incubated at 37°C with 5% CO_2_ for 5 h. The medium was then replaced with RPMI containing penicillin (100 µg/mL), streptomycin (100 µg/mL), and gentamicin (100 µg/mL), then incubated further for 2 h. Monolayers were washed and 100 µL of MTT-containing RPMI was added to each well, according to the kit protocol. After 2-4 h of incubation, 100 µL of solubilization reagent was added and ultimately A_570_ was measured to determine cellular respiration. RPMI with PBS alone served as the positive control and 0.4% Triton-X served as the negative control for cell viability.

### Hemolysis

Strains were cultured overnight in LB with aeration at 37°C. Overnight cultures were struck out for single colonies on tryptic soy agar containing 5% sheep blood and incubated overnight at 37°C. Plates were observed for zones of hemolysis surrounding the colonies.

### Hydrogen peroxide and acid resistance assays

Overnight bacterial cultures were incubated in LB medium at 37°C. OD_600_ was taken and cultures were normalized to 10^8^ CFU/mL in 3 mL of either plain LB or LB containing fresh 0.2% H_2_O_2_. Samples were immediately vortexed and serially diluted in PBS to enumerate CFU/mL at time 0 via drip-plating onto LB agar. Samples were incubated on the benchtop and then vortexed and plated at each time point.

For acid resistance the cultures were also normalized to 10^8^ CFU/mL in 3 mL of LB buffered with MES to pH 7, 5, or 2.5, and the input CFU/mL was determined. Cultures were then incubated for 2 h at 37°C with aeration and the final CFU/mL calculated.

### Serum resistance

Overnight LB cultures (1 mL) were pelleted by centrifugation and resuspended in 1 mL of sterile PBS. Suspensions were diluted 1:10 in sterile PBS. 10 µL of the diluted PBS solution was added to 190 µL of either 100% human serum or 100% heat-inactivated human serum. The bacterial inoculum was calculated by serially diluting and drip plating 20 µL of the bacteria-serum suspension on LB agar plates. The mix was then incubated for 90 minutes at 37°C, and then CFU was calculated in the same manner as the inoculum.

### Murine model of UTI

Female CBA/J mice (6-8 weeks old) were each transurethrally inoculated with 50 µL of a bacterial suspension (2×10^8^ CFU/mL) of each strain tested, using a sterile polyethylene catheter (I.D. 0.28 mm x O.D. 0.61 mm) connected to an infusion pump (Harvard Apparatus) as described (66, 67). The procedure was performed under anesthesia with ketamine/xylazine. Forty-eight hours post-inoculation, urine was collected and then mice were euthanized. Bladder, kidneys, and spleens were aseptically removed and homogenized. Urine and organ homogenates were serially diluted and plated onto LB agar to determine bacterial burden.

### Multiple linear regression and Pearson correlation

The averaged values of each assay (**Supplemental Table 2**) were loaded into RStudio (Version 1.41717). To calculate multiple linear regression, the baseR function lm() was used with three selected variables. To create a correlation matrix, the baseR function cor() was used with Pearson selected as the method.

## Acknowledgments

Financial support was provided by NIH R01AI165582 & R01AI059722 and NIAID F32AI147527.

We’d like to thank Stephanie Himpsl for her technical expertise and her help in editing the manuscript. We greatly appreciate Dr. Melanie Pearson for her intellectual conversation and manuscript editing.

## Figure Legends

**Supplemental Figure 1**. Growth of recent UPEC isolates, CFT073 and K12 in LB (A), M9 minimal medium with casamino acids (CAA) (B), glucose (C), glycerol (D), galactose (E), or ribose (F) as a sole carbon source, and pooled *ex vivo* human urine (G) Growth curves show averages of six biological replicates, error bars are ±SEM. (H) Indicated strains were cultured overnight in LB, and 5 µl of culture was spotted on CAS agar. The color change from blue to orange indicates iron chelation presumably due to siderophore production. An orange halo around the colony is due to diffusion of secreted siderophores.

**Supplemental Figure 2**. (A) Representative images of swimming motility assays of indicated strains. (B) Biofilm formation of clinical UPEC isolates, CFT073 and K12 in pooled *ex vivo* human urine at either 30°C or 37°C. Biofilms were stained with crystal violet and quantified through absorbance at 570 nm. Bars are an average of four biological replicates, error bars are ±SEM.

**Supplemental Figure 3**. (A) Representative image of an agarose gel displaying the result of an invertible element PCR assay. Bands representing *fimS* in the OFF or ON positions are labeled. (B) Representative image of a hemagglutination assay. The bottom row shows the addition of 1% mannose reverses the phenotype seen above, indicating that type 1 fimbriae are the adhesin responsible for the phenotype.

**Supplemental Figure 4**. Indicated strains were cultured overnight in LB, washed in PBS, then diluted to 10^8^ CFU/mL and incubated in heat-inactivated human serum for 90 minutes at 37°C. Input is enumerated in the black bars, while output is in the grey bars. Bars represent the average of four biological replicates, effort bars are ±SEM.

**Supplemental Figure 5**. Bacterial burden in the spleen of mice transurethrally inoculated with indicated strains. Mice were inoculated with 10^8^ CFU, and the infection progressed for 48 hours. Spleens were aseptically removed, homogenized, and the bacterial burden enumerated. Bars represent the median, dots represent individual animals.

**Supplemental Figure 6**. Pearson correlation matrix of all 18 phenotype assays. Values of used for each assay are shown in **Supplemental Table 2**. Asterisk denotes an R^2^ value greater than 0.5.

**Table S3.** Strains and plasmids used in this study.

| **<u>Strains</u>** | **<u>Description</u>** | **<u>Reference</u>** |
| --- | --- | --- |
| CFT073 | Wild-type pyelonephritis isolate (O6:K2:H1) | (65) |
| CFT073 L-ON | CFT073 ΔIRL, *fim* invertible element locked on | (68) |
| CFT073 L-OFF | CFT073 ΔIRL, *fim* invertible element locked off | (68) |
| K12 MG1655 | Non-pathogenic laboratory strain (OR:H48:K-) | (69) |
| HM1 | Wild-type cystitis strain isolated from a healthy young woman in 2009 | (31) |
| HM3 | Wild-type cystitis strain isolated from a healthy young woman in 2009 | (31) |
| HM6 | Wild-type cystitis strain isolated from a healthy young woman in 2009 | (31) |
| HM7 | Wild-type cystitis strain isolated from a healthy young woman in 2009 | (31) |
| HM14 | Wild-type cystitis strain isolated from a healthy young woman in 2009 | (31) |
| HM17 | Wild-type cystitis strain isolated from a healthy young woman in 2009 | (31) |
| HM43 | Wild-type cystitis strain isolated from a healthy young woman in 2009 | (31) |
| HM54 | Wild-type cystitis strain isolated from a healthy young woman in 2009 | (31) |
| HM56 | Wild-type cystitis strain isolated from a healthy young woman in 2009 | (31) |
| HM57 | Wild-type cystitis strain isolated from a healthy young woman in 2009 | (31) |
| HM66 | Wild-type cystitis strain isolated from a healthy young woman in 2009 | (31) |
| HM68 | Wild-type cystitis strain isolated from a healthy young woman in 2009 | (31) |
| HM86 | Wild-type cystitis strain isolated from a healthy young woman in 2009 | (31) |

**Table S4.**
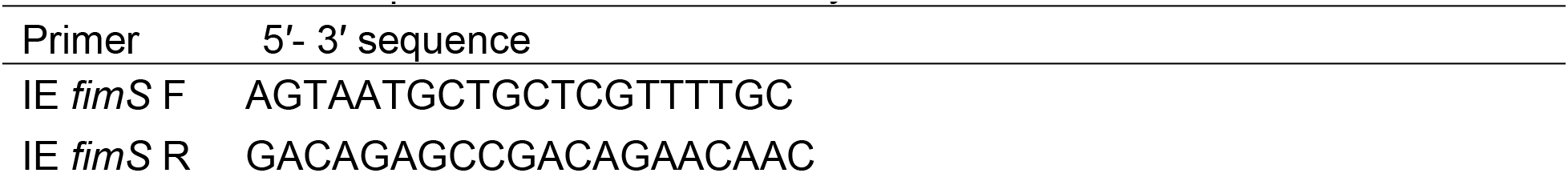
Primer sequences used in this study.

